# Predicting clinically promising therapeutic hypotheses using tensor factorization

**DOI:** 10.1101/272740

**Authors:** Jin Yao, Mark R. Hurle, Matthew R. Nelson, Pankaj Agarwal

## Abstract

Determining which target to pursue is a challenging and error-prone first step in developing a therapeutic treatment for a disease, where missteps are potentially very costly given the long-time frames and high expenses of drug development. We identified examples of successes and failures of target-indication pairs in clinical trials across 875 targets and 574 disease indications to build a gold-standard data set of 6,140 known clinical outcomes. We used information from Open Targets and others databases that covered 17 different sources of evidence for target-indication association and represented the data as a matrix of 21,437×2,211×17 with over two million non-null values. We designed and executed three benchmarking strategies to examine the performance of multiple machine learning models: Logistic Regression, Elasticnet, Random Forest, Tensor Factorization and Gradient Boosting Machine. With ten-fold cross validation, tensor factorization achieved AUROC=0.82±0.02 and AUPRC=0.71±0.03. Across multiple validation schemes, this was comparable or better than other methods. Tensor factorization is a general form of matrix factorization that has been successfully exploited in recommendation systems that suggest items to users based on their existing preference on a small number of items. Our application, using Bayesian probabilistic modelling, extends the capacity of matrix factorization to model multiple relationships between and among targets and indications. We use the model to show that our predicted probabilities of success correlate with clinical phases, and within clinical phase we can predict which trials are most likely to succeed.

## BACKGROUND

Drug discovery and development often begins with a drug target, typically a biological protein through which the drug exerts its therapeutic effect in patients. Although selecting an efficacious drug target is the first and most important step in drug development, more than half of clinical trials still fail due to lack of efficacy [1, 2]. This is critical to improve given the long-time frame and high expense of drug development. Major challenges in determining which targets to pursue and for which disease indication are that there are few positive examples (as most diseases do not have great proven treatments) and the evidence linking targets with disease mechanism is limited. Thus, any insights gleaned from the limited number of pursued targets may be useful in delivering new medicines with lower attrition rates. In this paper, we collated historical outcomes of clinical trials and determined if these clinical outcomes can be predicted retrospectively using multiple machine learning models built on existing evidence of the targets’ biological association with indications. A successful prediction model provides understanding on how informative the evidence is for clinical success, and is also capable of generating new target-indication hypotheses with higher potential of being developed into successful medicines.

One challenge in building such a model is that not all biological evidence is available for every pair of target and indication due to reasons such as technological limitation and limited disease coverage. For example, Open Targets is a recently published comprehensive database of molecular and phenotypical evidence that associates potential drug targets with disease indications [3]. As of June 2017, it contains 26,122 targets, 9,150 diseases with 2,857,732 positive associations from 15 data sources. Though Open Targets contains over 2.8 million associations, that is still only 0.08% of the possible combinations covered by this data, suggesting that a great deal of association evidence (99.92%) is still to be determined by biomedical researchers and clinicians. Traditional paradigms of machine learning algorithms, learning a mapping from input features (biological evidence) to output prediction (clinical outcomes), may be inadequate in this context. We show in this manuscript that the tensor factorization technique is useful in the analysis of this sparse biological dataset.

Tensor extends the concept of matrix to multidimensional array where each dimension corresponds to one “axis”, called mode, of a tensor [4]. Data in many applications can be naturally organized into a tensor format. **Figure 1a** shows a three-mode tensor representing different types of evidence associating targets with disease indications where the front “slice” represents clinical outcomes. Tensor factorization decomposes a tensor into factor matrices that compactly store information encoded in a tensor and integrate interaction across different modes even when a large portion of entries of a tensor is missing [4]. This technique has a wide range of applications such as in recommendation systems [5], knowledge graph systems [6] and multiple biomedical domains [7].

**Figure 1.**
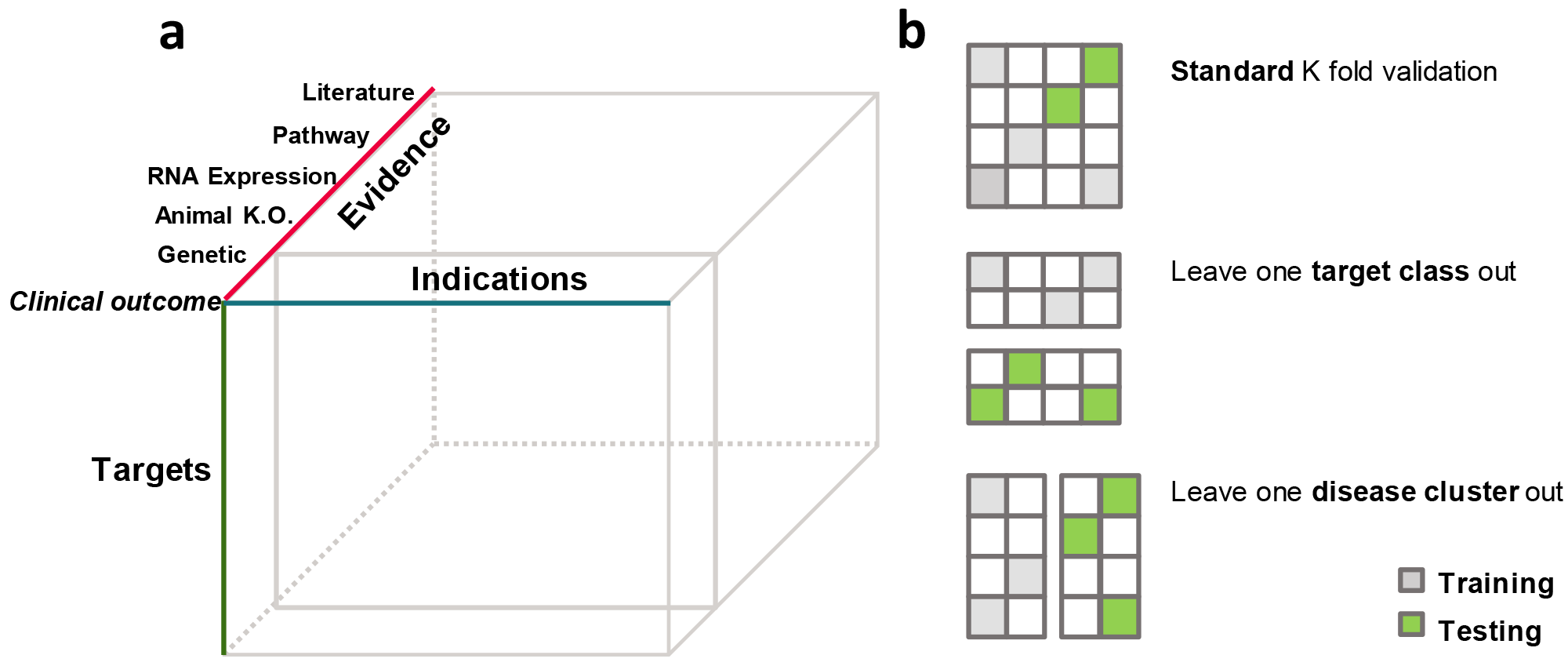
Data representation and model benchmark schematic. (a) Tensor representation of the dataset. The front “slice” matrix represents clinical outcomes of target-indication pairs. (b) Illustration of three schemes of benchmarking models on predicting clinical outcomes. Each matrix represents clinical outcomes of targets (rows) and indications (columns). Grey and green cells are target-indications pairs used for training and testing, respectively. Blank cells represent unknown clinical outcomes of target-indication pairs.

In the following sections, we first introduce the basic formulation of matrix factorization, a special form of tensor factorization. Then we explain our selection of a specific algorithm of tensor factorization based on characteristics of our problem. We then discuss how we design experiments to benchmark the method against a series of baseline models under three scenarios of drug discovery. We demonstrate that the model can capture known biological mechanisms of human diseases and can identify opportunities of approved drug targets to novel indications.

## METHODS

### Matrix factorization

The clinical outcomes of existing target-indication pairs can be represented in a matrix format as ***R*** ∈ ℝ^***M***×***N***^, where the M rows represent targets and the N columns represent indications. ***R***_***ij***_ = 1 if there is at least one drug that modulates target *i* and is marketed for indication *j*. ***R***_***ij***_ = 0 if all the drugs modulating target *i* are reported failed for indication *j* in the clinic (from Phase I to Phase III). For target-indication pairs that have no outcomes in the clinic, the corresponding ***R***_***ij***_ is empty. The goal is to predict clinical outcomes of all possible pairs of targets and indications i.e. fill out the empty ***R***_***ij***_′s. Thus, we can treat the problem as completing the target-indication matrix of clinical outcomes. Matrix completion problem has been widely studied in machine learning community in the context of recommendation systems [5, 8]. A famous application is Netflix’s movie recommendation system, where each user has ratings on a small number of movies and the task is to recommend movies for each user based on existing ratings of other users with similar patterns of movie ratings. Matrix factorization is recognized as one of more successful methods for this task [5, 9, 10]. The method assumes that the true completed matrix is of low rank and can be approximated by a product of two low-dimensional latent factor matrices that represent rows and columns of a matrix in a joint D-dimensional latent space, i.e. ***R*** ≈ ***U***^*T*^***V***, where 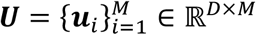, 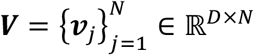 and **u**_***i***_ ∈ ℝ^*D*^, ***v***_*j*_ ∈ ℝ^*D*^ are column vectors of ***U*** and ***V***, respectively. The predicted entries in ***R***_***ij***_ is achieved by the inner product of ***u***_*i*_ and ***v***_***j***_. Learning of ***U*** and ***V*** can be formulated as an optimization problem by minimizing the mean squared error between observed and predicted entries. To avoid overfitting, regularization on the latent factor matrices is added to the minimization problem that can be solved by methods such as stochastic gradient descent and alternating least square [5].

### Bayesian tensor factorization

Many matrix-factorization based methods have been proposed for recommendation systems. To choose an appropriate method to predict clinical outcomes, we considered three aspects of our problem. First, some of the evidence is target-indication specific such as human genetic evidence for each disease, and this has been suggested as related to clinical outcome [11]. Second, in our data, there are several target-only attributes independent of indications, such as target protein location, gene expression in tissues etc. Thus, the chosen method should also take target-only information into consideration. Third, in drug discovery, it is not uncommon that targets or indications that have never been tested in clinical trials. In the case of movie recommendation systems, this corresponds to recommending movies to users who have not rated any movies in the system or recommending new movies that do not have any ratings in the system. The chosen method should be able to handle this situation.

Given these three aspects, we investigated a method based on tensor factorization, called Macau, that is capable of naturally handling all the three aspects in a unified Bayesian framework and was originally used to predict drug-protein interaction[12]. Tensor extends the concept of matrix to a multidimensional array, where each dimension corresponds to one mode of a tensor. Our data can be organized into a three-mode tensor: target × indication × evidence ***R*** ∈ ℝ^***M***×***N***×***K***^, where one entry ***R***_***ijk***_ indicates the association score in *k*^*th*^ evidence between target *i* and indication *j* and the clinical outcome matrix is included as one evidence; “slice” of the tensor (**Figure 1a**). Similar with matrix factorization, tensor factorization decomposes a tensor into a series of low-dimensional latent factor matrices where each matrix represents one mode of the tensor. One direct way to decompose a three-mode tensor is to assume that each entry ***R***_***ijk***_ can be expressed as the sum of elementwise product of three low dimensional vectors: ***u***_*i*_, ***v***_***j***_.and ***e***_***k***_, representing *i*^*th*^ target, *j*^*th*^ indication and *k*^*th*^ evidence respectively in a joint latent factor space. The prediction of clinical outcome is achieved by the sum of elementwise product of the low-dimensional representation of target, indication and clinical outcome evidence.

The specified tensor factorization method we chose is based on Bayesian probabilistic modeling, which assumes each evidence score is a random variable following a normal distribution. In this model, the parameter of the normal distribution is determined by the three low-dimensional latent factors: ***u***_*i*_, ***v***_*j*_.and ***e***_*k*_. Each latent factor is assumed to have a Gaussian prior with a Gaussian-Wishart hyper prior placed on its hyperparameters. The target-only attributes are linearly projected into the latent space and added to the target’s latent factor, which provides prediction ability to targets that do not have any observed clinical outcomes. The inference of model parameters and hyperparameters is carried out by Markov Chain Monte Carlo (MCMC) approximation method. Specifically, we used the Julia implementation of the method [13] and followed a common practice of MCMC inference where we “burn-in” samples generated in the beginning and collect samples after that to approximate posterior distribution over model parameters and hyperparameters [14]. In our case, the first 500 samples were discarded and the posterior distribution over parameters were estimated using 300 samples after the “burn-in” process. The predictive distribution is approximated from the 300 samples of the model parameters and then used to make predictions. Generally, we did not observe further improvement on prediction performance if we let the chain run longer. One parameter that needs to be specified is the number of latent factors. In this paper, we determined this parameter by a heuristic approach (supplementary material).

### Data collection and processing

We created a dataset which combined clinical outcomes from the commercial database Pharmaprojects [15] with evidence from Open Targets[3] and other sources (**Table 1**), converting all data to target-indication pairs using EntrezGene and MeSH ID ontologies to facilitate comparison. The collected evidence covers data space of 21,437 targets, 2,211 indications and 17 evidence sources. For outcome data, if at least one drug asset for a given target-indication pair was identified as successful, then the target-indication pair was classified as Succeeded. Of the remaining target-indication pairs, if at least one asset had a clinical failure then it was classified as a Clinical Failure. Open Targets presents evidence from each individual source as a numerical value for a target-indication pair, with a positive value representing evidence. To simplify the further collation of target-indication evidence with target-only evidence (**Table 2**), we converted numerical evidence value into binary values: 1 indicates positive association, 0 means that there is no association and unknown evidence is represented as null. We encoded categorical evidence, typically present as target annotations, as multiple binary values with each category converted into a binary value, i.e., having the property or not having the property. Here, we analyzed data mapped to the 574 non-cancer indications with at least one clinical outcome and the corresponding 875 targets. Oncology indications have different characteristics and the methodology can be expanded to those.

**Table 1.**
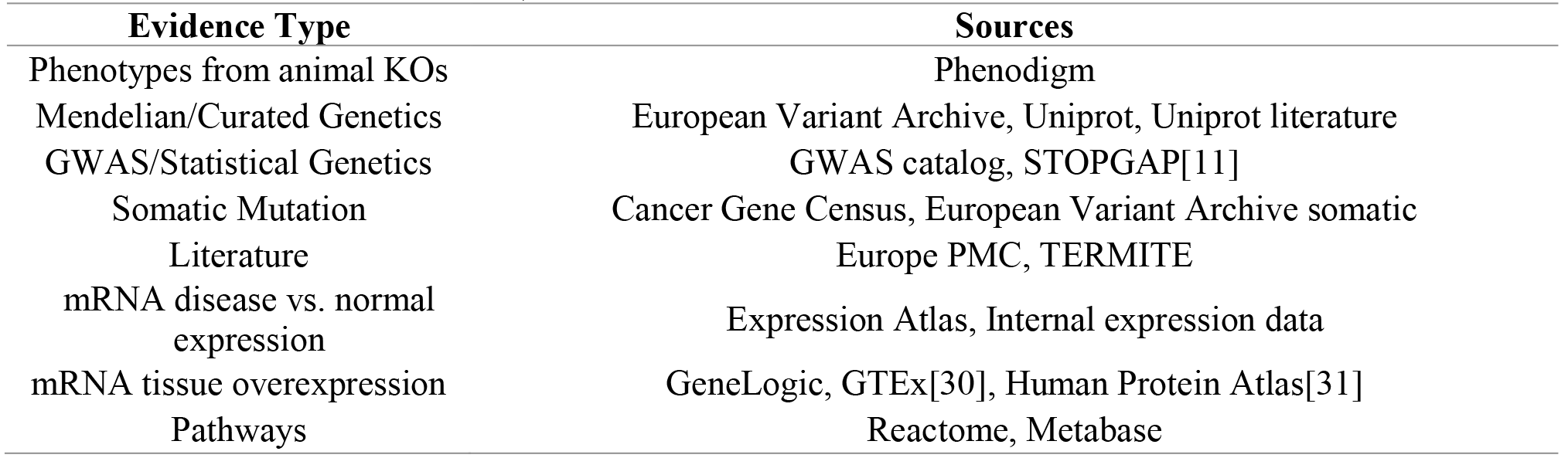
17 sources of target-indication evidence. Numerical data was obtained from Open Targets [1] except for commercial sources (TERMITE: www.scibite.com/products/termite; GeneLogic: GeneLogic Division, Ocimum Biosolutions, Inc) and where indicated

**Table 2.** Six sources of target-only evidence. Genes were broken into non-overlapping categories based on available data. Genes were classified as tolerant, intolerant and unclassified based on data from the Exome Aggregation Consortium[32] and the percentile rank of Residual Variation Intolerance Score [33]. Genes were based on identification of >=75% protein homology between human and mouse, data downloaded from BioMart[34]. Target Location and Topology were derived from a review of information from Gene Ontology, InterPro, PFAM and Uniprot,

| Evidence Type | Sources (# of categories) |
| --- | --- |
| Mutation Tolerance | ExAc_LoF(3), ExAc_Missense (3), RVIS (3), Mouse Protein identity (2) |
| Other Target Characteristics | Target Location (7), Target Topology (5) |

### Model benchmark experiments

We performed three cross validation experiments to evaluate the method in three scenarios (**Figure 1b**). In each experiment, we divided the target indication pairs with clinical outcomes (6,140) into experiment-dependent K folds, and tested the prediction results on a held-out fold using a model trained with the rest of folds. This process is repeated until every fold has been used as a test fold. The first case is a standard ten-fold cross validation where each fold is randomly determined but retains the same fraction of successes. In drug discovery, we know that certain sub-classes of targets and indications have different properties. In order to assess if the model can be generalized to sub-classes of targets and indications different from those used in the training stage, we devised two other cross validation experiments where each time the clinical outcomes of one pre-defined group of targets (indications) are left out as the test set. Specifically, for the leave one target group out case, we used the grouping defined by target class. A given target is assigned to one of ten target classes based on the target’s protein family retrieved from ChEMBL hierarchical target classification system [16]. For leave one indication group out case, we defined eight indication groups by *de novo* clustering indications based on how similar two indications are in terms of their relative positions in MeSH (Medical Subject Heading) hierarchical tree and co-occurrence frequency in literature (supplementary material).

### Baseline models

For comparison purposes, we also ran the cross-validation experiments using four machine learning models as baseline models. For the baseline models, each target indication pair is treated as a sample and the corresponding association evidence and target attributes are treated as its features. The task is being cast as a binary classification problem. To allow the four baseline models to handle missing values, we treated the association scores as categorical variables with three categories: no association (0), positive association (1) and unknown (missing) association. Each categorical variable is then encoded as two binary variables (also called one-hot encoding). The four models that we tested are:

1. Logistic Regression (LR), a simple linear model.
2. Elasticnet [17], which is a generalized linear model with elastic net regularization implemented in the glmnet R package where the regularization parameters were determined using cross validation.
3. Random Forest (RF), an ensemble model of decision trees where the parameters are determined by the Out-of-Bag error estimate using *tuneRF* function in *randomForest* R package.
4. Gradient Boosting Machine (GBM)[18], a boosting method which is implemented in *xgboost* [19] where we tuned the following parameters using cross validation: the feature shrinkage rate, maximum depth of a tree, subsample ratio of features and number of iterations.
5. Matrix Factorization (MF): We also included another baseline model where we *only* used the clinical outcome matrix and applied matrix factorization to complete the matrix for prediction. Specifically, we used a nuclear norm regularized matrix factorization method that is implemented in the *softimpute* [20] R package and the regularization parameter is determined through cross validation.

### Performance metrics

We used two metrics to measure the prediction performance of the evaluated methods: area under receiver operator curve (AUROC) and area under precision recall curve. (AUPRC). The AUROC measures the probability of a model ranking a randomly chosen positive example higher than a randomly chosen negative example and is commonly used in assessing performance of models for binary classification tasks. AUROC treats positive and negative examples equally, this metric is of limited value when the number of positive examples are relatively low. Given the low success rate in drug development, we chose AUPRC as the primary evaluation metric as it focuses on the performance on positive examples. Here the precision is the proportion of correctly predicted positives out of all predicted positives and recall is the proportion of correctly predicted positives out of all positives.

## RESULTS

### Model benchmark results

We performed a standard cross validation experiment to benchmark various types of machine learning models (**Figure 1b**, **panel 1**). The best model is the matrix factorization (MF) model (AUROC=0.83±0.02, AUPRC=0.77±0.02) (**Figure 2**), which only factorizes the clinical outcomes matrix without considering any other evidence in the dataset. Due to the highly-correlated structure within the clinical outcomes of target-indication pairs, the standard way of randomly splitting them into training and test sets may overestimate the predictability of clinical outcomes. This may explain the high performance of MF; knowing which targets have succeeded against which indications in the training data may provide enough information to predict the outcome status of new indications for these targets. Many drug targets are from the same gene family and it is very likely that targets within the same gene family are not assigned to the same training or test set, though the same drug may bind to members in each set. A similar effect may relate indications with different subtypes, as drug targets are often sequentially tried against closely related diseases.

**Figure 2.**
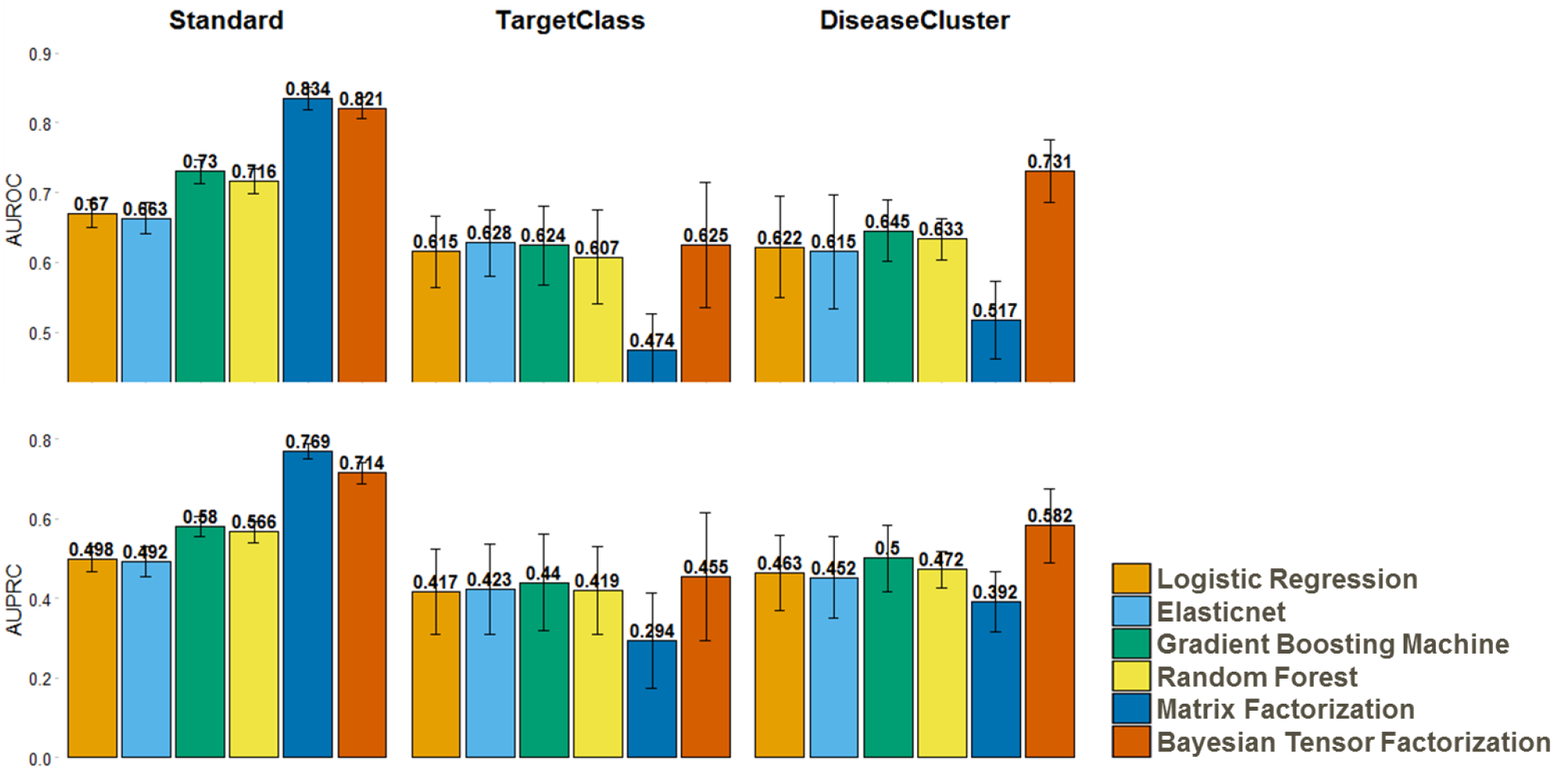
Benchmark performance of models. Prediction performance comparison in three benchmark schemes in terms of Area Under Receiving Operation Curve (AUROC, Top) and Area Under Precision Recall Curve (AUPRC, Bottom). Error bars are calculated from cross validation.

To mitigate this problem and obtain an unbiased estimation of predictive capacity, we designed two benchmark experiments, where a group of similar targets (**Figure 1b**, **panels 2 and 3**) or indications is held out as a test set and models trained on the other target or indication groups, respectively, are evaluated against the held-out set. We categorized targets into ten target classes largely derived from the ChEMBL hierarchical target classification system [16], and grouped indications into eight clusters that are based on MeSH hierarchy and co-occurrence frequency in literature (supplementary material). In the leave one target class out cross validation experiment (**Figure 2**), the performance of MF decreases dramatically as there is no information in the training set to predict clinical outcomes of the held-out target class. All the other methods perform similarly and the overall performance is not as good as in the standard cross validation setting. This implies that it is difficult to predict candidate indications for targets that have not been assessed in clinic trials. In the leave one disease cluster out validation experiment, the performance of MF again dropped below that of the other methods as there is no information about clinical outcomes of the held-out disease clusters in the training step.

However, the Bayesian tensor factorization (BTF) model scored as the best model in the disease group benchmark (AUROC=0.73±0.05, AUPRC=0.58±0.09) and the second best model in standard cross-validation (AUROC=0.82±0.02, AUPRC=0.71±0.03). It is counter-intuitive that BTF does not out-perform the MF method in the standard cross-validation case, as it incorporated more data. MF approach may be taking maximum advantage of the highly-related nature of the outcomes, given the poor performance of MF in the target class and disease group benchmarks. MF also only needs to learn latent factors to explain the clinical outcomes, while BTF needs to learn latent factors to explain the clinical outcomes and all the evidence as well, which is inherently a more difficult task.

In general, the performance of models that explicitly use evidence as predictors did not vary too much across three validation settings. Among these models, ensemble based methods (RF and GBM) worked slightly better than linear model based methods (LR and Elasticnet). Although MF performed relative well in the standard validation case, its performance was inconsistent among validation settings. BTF combined both evidence and inter-relationship among targets and indications and performed consistently well in all three validation scenarios.

### Leave one out experiment

One advantage of this leave one target/disease group out validation scheme is that we can also assess how trained models can be generalized to groups of targets/diseases that the models have never trained on before. Figure 3 shows the prediction performance of the six models on the held-out target classes (**Figure 3a**) and disease clusters (**Figure 3b**). In the leave one target class out case, the prediction performance averaged over the six models varies between target classes (AUPRC ranges from 0.24 to 0.58; AUROC ranges from 0.53 to 0.68). Specifically, we notice that the models perform consistently poorly for transcriptional factor targets and miscellaneous enzymes, which implies that these target classes are quite different from the other target classes. On the other hand, most models perform relatively well in protease targets. We note there is good consistency of performance among models within each target class, but this low variability is not repeated in the leave one disease cluster out case, where the prediction performance shows higher variability among disease clusters. For example, the BTF model performs better than the other models in the metabolic, GI and urologic and oral disease clusters, and performs as well as any other model in the other disease clusters.

**Figure 3.**
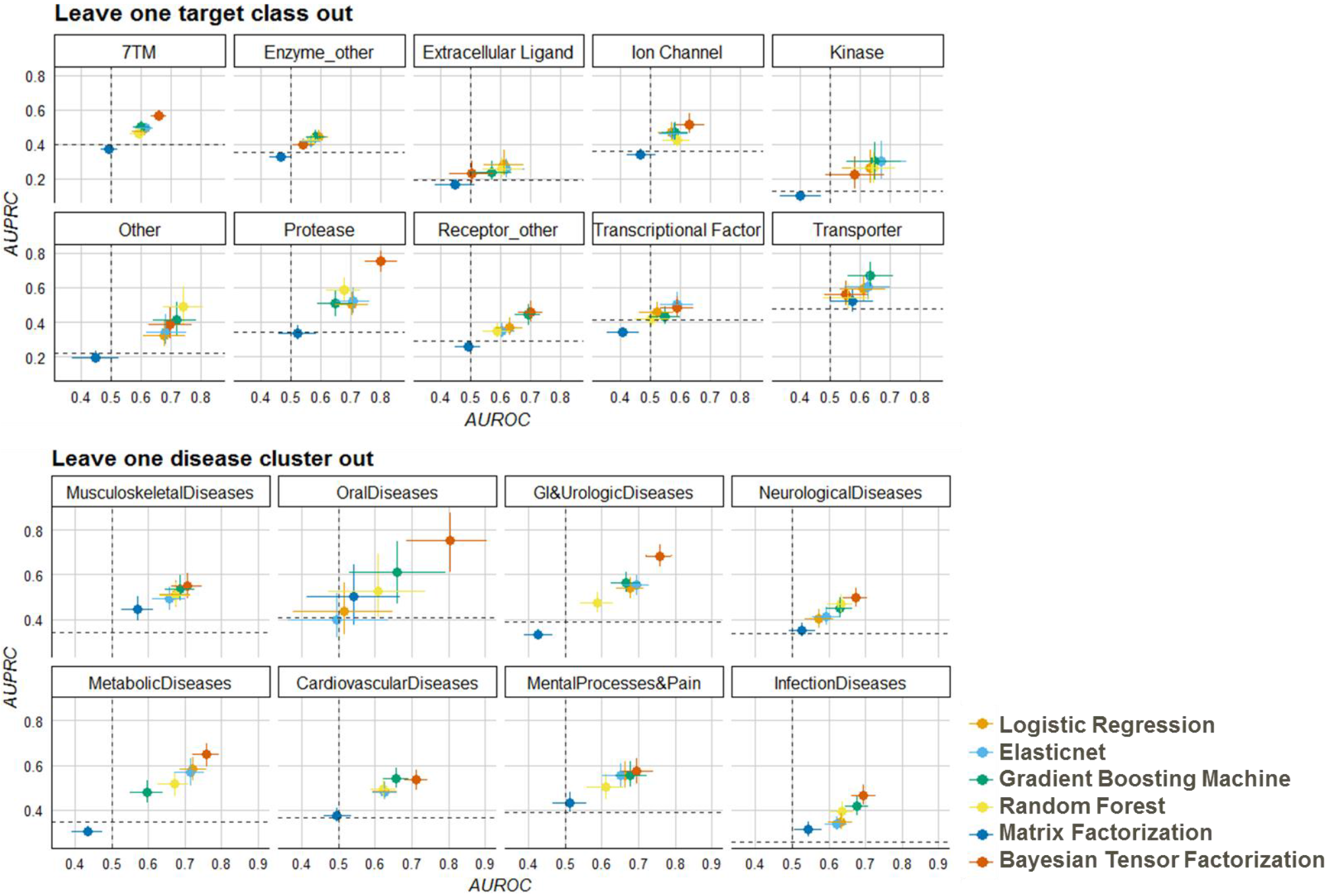
Benchmark performance of leave one out experiments. Model performance on predicting clinical outcomes of target classes (Top) and disease clusters (Bottom) in leave one out experiment in terms of Area Under Receiving Operation Curve (AUROC, x axis) and Area Under Precision Recall Curve (AUPRC, y-axis). 95 confidence interval are calculated using 1,000 bootstraps. Dotted lines mark the AUROC (vertical) and AUPRC (horizontal) of random guess, which is 0.5 and% of positives in the testing set, respectively.

### Latent factors capture disease relationship

After benchmarking the performance of the BTF model in the cross-validation experiments, we fitted the model to the whole dataset. We chose eleven latent factors (supplementary material). Before using the fitted BTF model to make any predictions, we explored whether the latent factors learned from the model are biologically meaningful so that we can increase our trust on the prediction made by the model. To do so, we reduced the eleven latent factors to two dimensions using t-SNE [21] to visualize how indications are distributed and examined whether the learned latent factors can capture inter-relationship among indications. t-SNE is a dimension reduction technique used to visualize highdimensional dataset where similar points in high dimensional space are transformed to neighboring points in a low dimensional space and dissimilar points are transformed to distant points in the low dimensional embedding. **Figure 4a** shows the two-dimensional t-SNE embedding of the 574 indications with at least one clinical outcome, where large-scale themes as well as local clustering of biological mechanisms can be observed. For example, different disease areas generally occupy different domains in the map. Three clusters are enriched with three distinct disease categories including Central nervous system diseases, Digestive system diseases and Hemic & lymphatic diseases, respectively. Auto-immune diseases, such as rheumatoid arthritis, asthma, psoriasis and Crohn’s diseases, that manifest in different organs are localized in the same neighboring area in the map. Figure 4a shows the 2D embedding with perplexity=30 in t-SNE. The above observation is consistent using different perplexity values in the range from 10 to 50.

**Figure 4.**
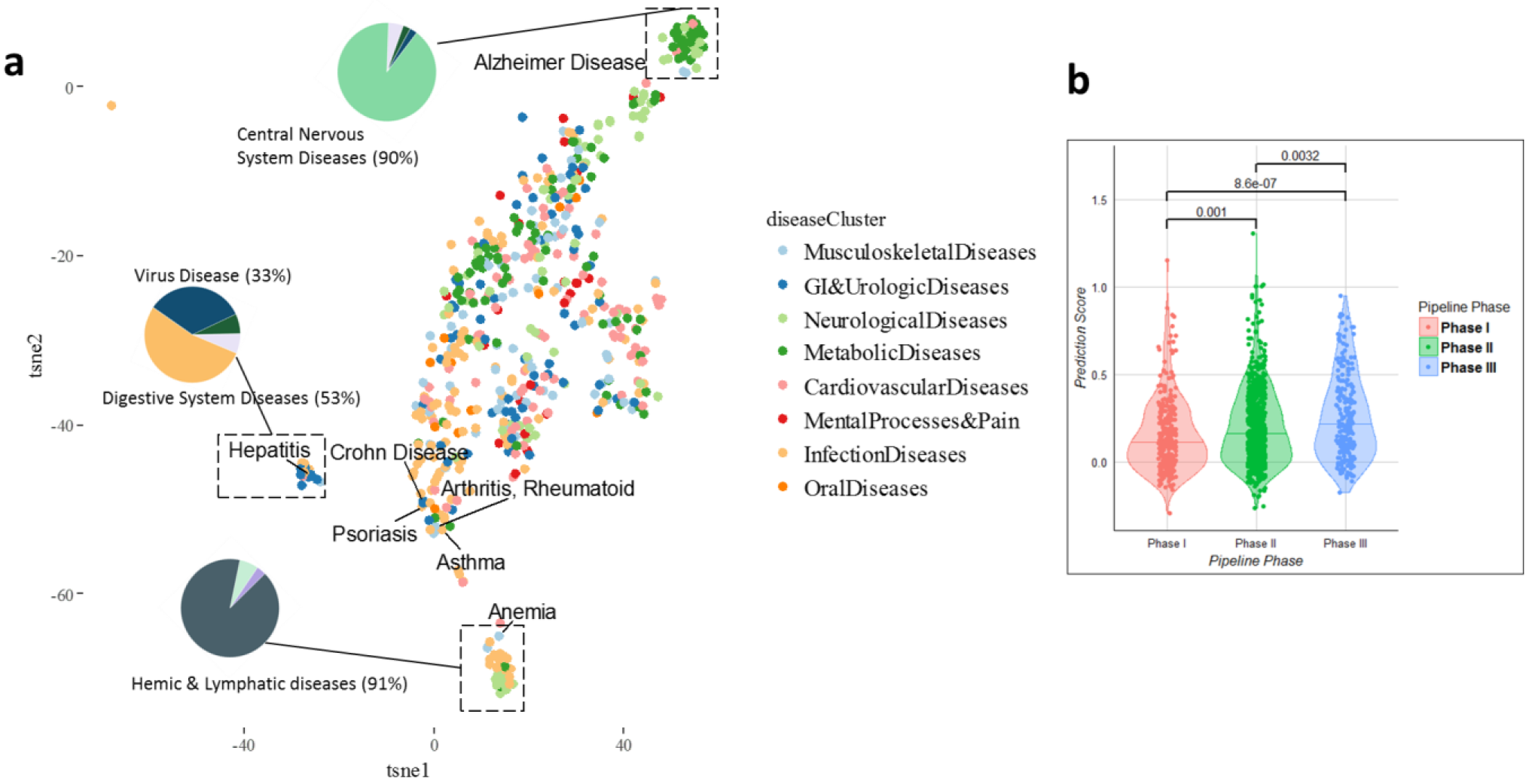
Validation of BTF model prediction. a). t-SNE visualization of indications based on the latent factors learned in BTF model. Each dot represents one indication and the size of the dot is proportional to the number of targets that have been clinically failed. The inserted pie charts show diseases composition of representative clusters of indications in the 2D visualization. b) BTF prediction scores of target-indication pairs in Phase I-III clinical trials. P-values are based on Wilcoxon rank sum tests.

### Prediction scores of target-indication pairs under clinical trials

To validate the prediction made by the BTF model, we chose 1,246 novel target-indication pairs that were in progress in clinical trials (Phase I-III) at the time when we collected the data (May 2016), and thus did not have clinical outcome readouts. We compared the prediction scores generated by the BTF model on these target-indication pairs and noticed that the prediction scores of later phase pairs are significantly higher than those of earlier phase pairs (**Figure 4b**), which recapitulates the observation that drugs in later phases on average have higher likelihood of approval. Since we did not include phase information of these target-indication pairs when training the model, these pairs serve as an independent test set and the results increase our confidence on the predictions of the model.

Next, we conducted literature search on the top 63 hypotheses of the 1,246 pairs based on a prediction score threshold, which corresponds with 0.8 precision and 0.27 recall in the standard cross validation experiment. We list 15 of these 63 hypotheses along with a relevant literature reference in Table 3; the complete list of 63 can be found in the supplementary material.

**Table 3.** High Scoring Pairs of Interest from TF Model. New indications of approved targets in clinical trials (Phase* as of May 27, 2016) that have the highest probability of eventual clinical success as measured by the tensor factorization model. The full list is available in the supplement. For illustrative purposes, we list a related indication approved for assets for each target

| Target | High Scoring Indication in Clinical Pipeline (Phase*) | PubmedID | Related Approved Indication (Total Approved Indications) |
| --- | --- | --- | --- |
| ABCC8 | Glucose Intolerance (III) | 23903354 | Diabetes Mellitus (1) |
| ADRB1 | Cachexia (II) | 20426789 | Ischemia (13) |
| ADRB2 | Hypoglycemia (I) | 22013013 | Glaucoma (15) |
| ADRB2 | Myocardial Infarction (III) | 26692153 | Heart Failure (15) |
| AGTR1 | Hypercholesterolemia (III) | 12117739 | Hyperlipidemias (7) |
| CYP3A4 | Hepatitis C (II) | 20938912 | HIV Infections (1) |
| IL2 | Behcet Syndrome (II) | 26654556 | Graft Rejection (1) |
| IL6 | Waldenstrom Macroglobulinemia (I) | 26238488 | Giant Lymph Node Hyperplasia (1) |
| IL6 | Arthritis, Psoriatic (II) | 27789987 | Giant Lymph Node Hyperplasia (1) |
| OPRM1 | Schizophrenia (III) | 27397309 | Migraine Disorders (22) |
| RYR1 | Muscular Dystrophy, Duchenne (I) | 26793121 | Malignant Hyperthermia (1) |
| SERPINC1 | Hemophilia (II) | 27099538 | Blood Coagulation Disorders (16) |
| TNFSF11 | Hypercalcemia (II) | 27904108 | Osteoporosis (1) |
| VDR | Alopecia (I) | 27932380 | Keratosis (9) |
| VDR | Cachexia (I) | 22497530 | Chronic Kidney Failure (9) |

As an example, interleukin 6 (IL6) is an approved drug target for giant lymph node hyperplasia (**Table 3**). Our results suggest that the current trials for psoriatic arthritis, which includes a Phase IIb trial of a monoclonal antibody against this protein [22], have greater than random chance of success.

Psoriatic arthritis is a chronic inflammatory arthritis that is associated with psoriasis and thus somewhat related to the successful indication for IL6, a cytokine with a wide variety of biological functions. It induces the acute phase response and is involved in the final differentiation of B-cells into Ig-secreting cells in lymphocyte and monocyte differentiation. It acts on B-cells, T-cells, hepatocytes, hematopoietic progenitor cells and is required for the generation of T(H)17 cells. It also acts as a myokine and is discharged into the bloodstream after muscle contraction and acts to increase the breakdown of fats and to improve insulin resistance[23]. Genetic polymorphism of IL6 has been shown to be significantly associated with a form of psoriatic arthritis [24], and serum IL6 is considered as a biomarker for assessing disease activity in patients with psoriasis, as well as for predicting responsiveness of joint symptoms to biologic treatment [25].

Another target of interest is angiotensin II receptor type 1 (AGTR1), an important effector controlling blood pressure and volume in the cardiovascular system. It has been approved for many cardiovascular indications such as heart failure, myocardial infarction and hypertension. The predicted indication for AGTR1 is hypercholesterolemia, also known as high cholesterol. AGTR1 antagonism improves hypercholesterolemia-associated endothelial dysfunction[26] and attenuates the inflammatory and thrombogenic responses to hypercholesterolemia in venules [27]. Significant association of AGTR1 polymorphism with hypercholesterolemia was also observed in hypertension patients [28]. During the time of preparing this manuscript, we found that a drug has been launched in South Korea [29] for hypercholesterolemia by targeting AGTR1.

## DISCUSSION

In this paper, we focused on the problem of predicting clinically promising therapeutic hypotheses using associative knowledge of targets and indications. We compared tensor factorization with other traditional machine learning methods in a variety of benchmarking experiments and identified two interesting findings from the evaluation of this method: 1) the latent factors learned from the model align with known biological relationships among human diseases; and 2) the method can be applied to different scenarios of drug discovery and achieves competitive prediction performance.

However, there are some limitations worth discussing before deploying tensor factorization to propose novel target-indication hypotheses. First, the model relies on the available compilation of evidence sources. Open Targets provided us a good foundation, but clearly more sources could be gathered. Second, we treated every clinical failure equally. Our preliminary analysis has shown that some target-indications pairs have been tried multiple times and are still being pursued clinically, while some failed only once and were never tested again. Although the probabilistic framework of the model can potentially mitigate this problem, the model does not explicitly differentiate definitive failures from those that have not been thoroughly explored and may become successful drugs in the future. Lastly, we only applied the technique to a dataset of targets and indications with at least one clinical outcome; thus, the application as benchmarked here is constrained to applying approved drug targets to new indications. The methodology, however, can be expanded to any target and any indication so long as their evidence is encoded in the data. Such an application may result in the identification of novel target-indication hypotheses with a high predicted probability of being successfully translated into medicines.

## CONCLUSION

In this work, we evaluated a machine learning technique called tensor factorization on the problem of predicting clinical outcomes of therapeutic hypotheses using existing association evidence between drug targets and disease indications. We illustrate that the method can achieve equal or better prediction performance compared with a variety of baseline models across three scenarios of drug discovery, and the learned model can capture known biological mechanism of human diseases. Furthermore, we demonstrated an application of the method to predict outcomes of trials on novel indications of approved drug targets. Future work includes expanding this method to targets and indications that previously have never been clinically tested and proposing novel target-indication hypotheses that can be developed into medicines with predicted high probabilities of success.

## ACKNOWLEDGMENTS

We would like to thank Andrew Rouillard for his helpful discussion and valuable comments on the manuscript. We thank Johannes Freudenberg for supplying the data for disease grouping. We thank Gautier Koscielny for facilitating access to Open Target data. We also thank Mandy Bergquist, Enrico Ferrero, Victor Neduva and Naruemon Pratanwanich for helpful discussions.

